# TM3’seq: a tagmentation-mediated 3’ sequencing approach for improving scalability of RNA-seq experiments

**DOI:** 10.1101/585810

**Authors:** Luisa F. Pallares, Serge Picard, Julien F. Ayroles

**Affiliations:** Lewis-Sigler Institute for Integrative Genomics, Princeton University, Princeton, USA; Department of Ecology and Evolution, Princeton University, Princeton, USA

**Keywords:** RNA-seq, Tn5 transposase, transcriptomics, library preparation, TM3seq

## Abstract

RNA-seq has become the standard tool for collecting genome-wide expression data in diverse fields, from quantitative genetics and medical genomics to ecology and developmental biology. However, RNA-seq library preparation is still prohibitive for many laboratories. Recently, the field of single-cell transcriptomics has reduced costs and increased throughput by adopting early barcoding and pooling of individual samples —producing a single final library containing all samples. In contrast, RNA-seq protocols where each sample is processed individually are significantly more expensive and lower throughput than single-cell approaches. Yet, many projects depend on individual library generation to preserve important samples or for follow-up re-sequencing experiments. Improving on currently available RNA-seq methods we have developed TM3’seq, a 3’-enriched library preparation protocol that uses Tn5 transposase and preserves sample identity at each step. TM3’seq is designed for high-throughput processing of individual samples (96 samples in 6h, with only 3h hands-on time) at a fraction of the cost of commercial kits ($1.5 per sample). The protocol was tested in a range of human and *Drosophila melanogaster* RNA samples, recovering transcriptomes of the same quality and reliability than the commercial NEBNext^®^ kit. We expect that the cost-and time-efficient features of TM3’seq make large-scale RNA-seq experiments more permissive for the entire scientific community.

## Introduction

The type of questions biologists ask depend on the interests and curiosity of each scientist. However, the type of questions we get to answer depend, to a great extent, on the experimental and analytical methods that we have at hand. RNA sequencing (RNA-seq), as a mean to assess genome-wide gene expression profiles, is one example of a method that has revolutionized research in biology. The possibility of collecting genome-wide expression data offers an unbiased way to integrate genotypic variation, phenotypic variation, and the environment; this has made RNA-seq a very popular tool in a wide range of biological disciplines, from ecology to developmental biology to quantitative genetics. However, RNA-seq remains prohibitive for many laboratories due to the high cost of library preparation using commercial kits.

In the last couple of years, some attempts have been made to reduce cost and increase the throughput of RNA-seq library preparation. Most of these protocols were developed to address the challenges in single-cell transcriptomics, where the RNA input is very low, and sample sizes easily reach thousands of cells (Macosko et al. 2015, Hashimshony et al. 2016). A general RNA-seq library preparation procedure consists of five main steps: reverse transcription (RT), cDNA synthesis, cDNA fragmentation, adapter ligation, and PCR amplification —the order in which such steps are performed varies depending on the specific protocol. Single-cell protocols adopted early barcoding of samples, that is, each sample is labeled with unique identifiers during RT, and therefore, individual samples can be pooled after RT and processed as a single sample for the rest of the protocol. Such early pooling (or early multiplexing) of samples reduces the costs per individual sample and increases throughput. Recently, BRB-seq successfully adapted the early multiplexing approach of single-cell protocols to bulk RNA-seq (Alpern et al. 2019). However, such feature comes with two main caveats: first, the identity of the individual samples is only recovered during bioinformatic analysis, and second, any variation in RNA input amount will result in high variation in sequencing coverage per sample (see the SMART-seq2 method for an exception (Picelli et al. 2014)). The early multiplexing feature is therefore not suitable for experimental designs where individual libraries are required for follow-up re-sequencing experiments or because the original samples are irreplaceable. Under those scenarios, it is desirable to use protocols that pool individual samples after final PCR amplification (i.e. late multiplexing).

Surprisingly, there have been far fewer attempts at reducing costs and increasing throughput of RNA-seq library preparation protocols while preserving the individuality of each sample at every step. From the field of ecological genetics, a 3’-enriched protocol – TagSeq – was proposed by Meyer et al. (2011) and optimized by Lohman et al. (2016). Their results showed that 3’-enriched libraries recover the transcriptional profile of individual samples with the same or better quality than libraries prepared with the commercial kit NEBNext^®^, with the caveat that TagSeq is not suitable for alternative splicing studies and does not recover information (e.g. polymorphism) from the entire coding sequence. By focusing on the 3’ end of mRNA molecules, fewer reads are required to cover the whole transcriptome compared with traditional full-transcript-length approaches, therefore reducing sequencing costs. A second approach, Smart-seq2, generates full-length RNA-seq libraries in a time-and cost-effective manner by using hyperactive Tn5 transposase which fragments the DNA and incorporates adapters in one step, therefore reducing hands-on time. The Smart-seq2 authors, as well as others (Picelli et al. 2014, Hennig et al. 2018), have made available protocols to produce home-made Tn5 transposase for a fraction of the cost of commercial Tn5, resulting in a substantial drop in the reagent cost of library preparation.

Here we present TM3’seq – Transposase Mediated 3’ RNAseq – a new late multiplexing RNA-seq protocol that builds on the 3’ approach of TagSeq (Lohman et al. 2016) and the Tn5-based library preparation of SMART-Seq2 (Picelli et al. 2014) to generate 3’-enriched mRNA libraries. It requires minimum hands-on time and is ∼23 times cheaper ($1.5 per sample) than the commercial gold-standard kit (NEBNext^®^Ultra, $35-$44per sample), while preserving the individuality of each samples at every step. To further reduce the sequencing costs, TM3’seq incorporates dual-index sample barcodes to allow high multiplexing, but in contrast with early multiplexing methods (e.g. BRB-seq) that require pair-end sequencing, TM3’seq barcodes can be read using single-end sequencing, further reducing costs

The technical and biological performance of TM3’seq was evaluated by processing human blood and adipose RNA and comparing the results to the commercial NEBNExt^®^ Ultra Directional RNA kit. In addition, to evaluate the performance of the TM3’seq method when processing large numbers of samples of low and variable RNA input, we generated transcriptome profiles — separately for head and body — of 48 *Drosophila melanogaster* flies. Our results show that TM3’seq reliably recovers genome-wide gene expression profiles of different tissue types in two different taxa, humans and flies. The full TM3’seq protocol, including the RNA extraction step, has been implemented in 96-well plates format to facilitate its implementation on liquid-handling robots. Finally, to facilitate the analysis of the data generated by TM3’seq, we have generated a straightforward analysis pipeline that will allow non-experts to go from the raw FASTQ files delivered by the sequencing facilities to the expression profile of each sample, generating output files that can be directly imported into widely-used software for gene expression analyses. The protocols and data analysis pipeline can be found on lufpa.github.io/TM3Seq-Pipeline.

## Methods

### 1. Samples

#### Human RNA

We used commercially available total RNA from human adipose tissue (Clontech, lot 1604416A) and blood (peripheral leukocytes) (Clontech, lot 1002007) to evaluate the performance of our library preparation protocol – TM3’seq – and compare it to the performance of a commercial kit commonly used to generate RNA-seq libraries – NEBNext^®^ Ultra™ Directional RNA kit (NEB, #E7420S). Three replicates of 200ng total RNA per tissue (blood and adipocyte) were used as input for NEBNext^®^ and TM3’seq library preparation methods. NEBNext^®^ libraries were prepared following the manual instructions for Poly-A mRNA isolation method. The TM3’seq protocol is described below, and the detailed step-by-step protocol as well as the list of oligonucleotides used are available as Suppl. File 1 and Suppl. Table 1, respectively. The protocol is also available on lufpa.github.io/TM3Seq-Pipeline.

#### Fly tissue

48 female *Drosophila melanogaster* flies were collected from an outbred population kept at 25°C with 65% humidity and a 12h/12h light/dark cycle. Flies were decapitated to separate heads and bodies. Each one of the 48 heads and 48 bodies was placed in a well of a 96-well plate. One 2.8mm stainless steel grinding bead (OPS diagnostics, #089-5000-11) and 100μl of lysis buffer (see Suppl. File 2 for details on this buffer) were added to each well containing heads or bodies. The tissue was homogenized for 10 minutes at maximum speed in a Talboys High Throughput Homogenizer (#930145). The resulting lysate was transferred to a new 96-well plate for mRNA isolation. TM3’seq uses oligo-dT-primed cDNA synthesis and therefore mRNA isolation is not necessary. However, we found that adopting bead-based mRNA isolation approaches was more straightforward, required less steps, and was cheaper than bead-based total RNA isolation. mRNA extraction was performed using Dynabeads™ mRNA DIRECT™ Purification Kit (ThermoFisher, #61012) following the protocol from Kumar et al. (2012) that we optimized for low input and low cost per individual sample. The detailed step-by-step protocol is available as Suppl. File 2 and in webpagelink. The mRNA yield was 10-20ng from a single head, and 90-180ng from a body, and therefore TM3’seq protocol was optimized to produce high quality libraries from small amounts of input material like single *Drosophila* heads.

After tissue homogenization, the processing of the samples (mRNA isolation and library preparation) was done in the CyBio^®^ FeliX liquid handling robot (Analitik Jena) to allow high-throughput while reducing the variability introduced by manual handling of individual samples. The detailed protocols in Suppl. File 1-2 can be used as reference for the implementation of sample processing in liquid-handling platforms. For the specific case of CyBio^®^ FeliX, the main protocols, subroutines, and instructions to set-up the robot are available upon.

### 2. TM3’seq protocol

The list of oligonucleotides used in this protocol is available as Suppl. Table 1, and details on buffers composition can be found in Suppl. File 1. *First strand cDNA synthesis*. Before reverse transcription, 200ng of total RNA in 10μl (or 10ng of mRNA in 10 μl) was mixed with 1μl Tn5ME-B-30T 0.83uM oligo and incubated at 65°C for 3 minutes. This oligo has both adapter-B complementary to the i7 primer that will be used to amplify the final libraries, and a poly-T sequence of 30nt that binds to the poly-A tail of mRNA molecules. The use of this oligo to prime the first strand cDNA synthesis results in libraries enriched for the 3’ end of mRNA. Reverse transcription was done by adding the following reagents to the reaction described above: 1μl SMARTScribe ™ RT (Takara, #639538), 1μl dNTPs 10mM (NEB, #N0447S), 2μl DTT 0.1M (Takara, #639538), 4μl 5× First-Strand buffer (Takara, #639538), and 1μl B-tag-sw oligo. This oligo allows for the template switching at the 5’ end of the mRNA molecule to incorporate a universal 3’ sequence during first strand cDNA synthesis. The biotin in this oligo helps prevents concatemerization of the oligo, a common problem when the input RNA amount is low. Synthesis of the first cDNA strand was done at 42°C for 1h, followed by 15min at 70°C to inactivate the reverse transcriptase.

#### cDNA amplification

5μl of the first-strand reaction was mixed with 7.5μl of OneTaq HS Quick-load 2× (NEB, #M0486L) and 2.5μl water and amplified for three PCR cycles following the program: 68°C 3min, 95°C 30sec, [95°C 10sec, 55°C 30sec, 68°C 3min] *3 cycles, 68°C 5min.

#### cDNA Tagmentation

10μl (100μM) forward oligo (adapter-A) and 10μl (100μM) reverse adapter A oligo (Tn5MErev) were mixed with 80μl re-association buffer (10mM Tris pH 8.0, 50mM NaCl, 1mM EDTA), and annealed in a thermocycler following the program: 95°C 10min, 90°C 1min, reduce temperature by 1°C/cycle for 60 cycles, hold at 4°C. The annealed adapter-A binds to the Tn5 transposase and forms the free-end adapters that will be ligated to the cDNA after the Tn5 transposase fragments the cDNA. To load the adapter-A into Tn5, 5 μl of annealed adapter-A (1μM) were mixed with 5μl of homemade Tn5 (made following Picelli et al. (2014)) and incubated in a thermal cycler for 30min at 37°C. The adapter-A sequence is complementary to the i5 primer that will be used to amplify the final libraries. Adapter-B was added in the first step of first strand synthesis. The pre-charged Tn5 was diluted 7× in re-association buffer:Glycerol (1:1). 5μl of cDNA was mixed with 1μl of pre-charged Tn5, 4μl of TAPS buffer 5× pH 8.5 (as described in Picelli et al. (2014); 50mM TAPS, 25mM MgCl_2_, 50% v/v DMF), and 5μl of water, and the solution was incubated for 7min at 55°C. 3.5μl of SDS 0.2% (Promega, #V6551) was added to the solution and incubated in a thermal cycler for 7min at 55°C to dissociate the Tn5 that remained bound to the cDNA.

#### Final library amplification

Finally, 10μl of OneTaq HS Quick-Load 2x (NEB, #M0486L), 1μl i5 primer 1uM, 1μl i7 primer 1μM, and 7μl of water were used to amplify 1μl of the tagmentation reaction following the program: 68°C 3min, 95°C 30sec, [95°C 10sec, 55°C 30sec, 68°C 30sec] *12 cycles, 68°C 5min. The number of amplification cycles ranged between 12 and 18 depending on the experiment and the desired library yield; it should be noted that the number of amplification cycles does not substantially affect the results (see Fig 3); the number of cycles used is specified in each figure legend.

#### Pooling of individual libraries and sequencing

Human RNA libraries were individually cleaned using Agencourt AMPure XP beads (Beckman Coulter, #A63881) and a double-sided procedure (left 1×-right 0.6×); after cleaning, all samples were pooled. Fly heads and bodies were cleaned in separate pools using AMPure XP beads as described above, and equal proportions of each pool were combined after cleaning. Human and fly samples were sequenced in independent runs on an Illumina HiSeq 2500, using dual indexes (Index1(i7) – 8bp, Index2(i5) – 8bp) and single-end 67bp sequencing. All sequencing was done at the Genomics Core Facility at the Lewis-Sigler Institute for Integrative Genomics at Princeton University.

### 3. Expression level quantification

RNA-seq reads were trimmed for low quality bases and adapter sequences using Trimmomatic 0.32 [parameters: *SE ILLUMINACLIP:1:30:7 LEADING:3 TRAILING:3 SLIDINGWINDOW:4:15 MINLEN:20*] (Bolger et al. 2014). Reads shorter than 20nt after trimming were discarded. Trimmed reads were mapped to the GRCh38 assembly of the human genome using STAR [--outSAMmapqUnique 60 – outSAMtype BAM SortedByCoordinate] (Dobin et al. 2013). Only uniquely mapped reads were kept for further analysis (*samtools view –q 50*) (Li et al. 2009). The resulting bam files were down sampled *(samtools view –s)* depending on the comparison being made. The number of reads per sample used in each analysis is stated in the figure legends.

For human samples, uniquely mapped reads were assigned to the set of 20612 annotated genes in the GRCh38 assembly of the human genome using featureCounts from the package Subread 1.5.1 [*featureCoutns -t exon –g gene_id*] (Liao et al. 2014). The raw read counts were imported into R (R-Core-Team 2018) and analyzed with DESeq2 (Love et al. 2014) to identify differentially expressed genes between blood and adipocytes (design = ∼ tissue) using the model *gene expression* ∼ *tissue*. P-values were adjusted for multiple testing using the Bonferroni method, and the threshold for significance was set to p-value< 0.05.

For fly samples, uniquely mapped reads were assigned to the set of 17472 annotated genes in the r6.14 assembly of the *Drosophila melanogaster* genome using feautureCounts from the package Subread [*featureCoutns -t exon –g gene_id*] (Liao et al. 2014). Only samples with more than 500k reads assigned to protein-coding genes were included in further analysis (n=74, 39 bodies, 33 heads). The raw read counts were imported into R (R-Core-Team 2018) and analyzed with DESeq2 (Love et al. 2014) to detect differentially expressed genes between head and body (design = ∼ body part). P-values were adjusted for multiple testing using the Bonferroni method, and the threshold for significance was set to p-value< 0.05.

### 4. ERCC probes

ERCC probes are a common set of polyadenylated RNA molecules designed by the External RNA Controls Consortium (ERCC) to be used as control RNA of known concentration that can be compared between different experiments. ERCC probes (Thermo Fisher, #4456739) were spiked into an independent set of samples that were processed with both protocols (TM3’seq and NEBNext^®^) for a total of two ERCC replicates per method. ERCC sample 1 corresponds to ERCC Mix 1, and sample 2 to ERCC Mix 2. Mix 1 and 2 only differ in the concentration of each of the probes, and therefore allow differential gene expression analyses.

Reads were mapped to the ERCC92.fa sequence file downloaded from Thermofisher.com using STAR (Dobin et al. 2013). The bam files from all samples were down sampled to 100k uniquely mapped reads *(samtools view –s)*, and these reads were assigned to the 23 ERCC probes using featureCounts (Liao et al. 2014). The raw counts were imported into R (R-Core-Team 2018) for further analyses performed with ERCC-Dashboard (Munro et al. 2014).

### 5. Pipeline for processing of RNA-seq FASTQ files

We have developed a pipeline that allows the processing of raw FASTQ files into gene counts files that are ready to be used as input in standard differential expression analysis software like DESeq2 (Love et al. 2014), or in eQTL mapping software like MatrixEQTL (Shabalin 2012). The pipeline is available on lufpa.github.io/TM3Seq-Pipeline.

### 6. Data availability

FASTQ files are available under the SRA accession PRJNA528324. mRNA extraction and TM3’seq protocols are available as Suppl. File 1 and Suppl. File 2 and in lufpa.github.io/TM3Seq-Pipeline.

## Results

Here we present TM3’seq –Transposase Mediated 3’ RNAseq–, a 3’-enriched RNA-seq protocol to generate high-quality libraries in a cost- and time-efficient manner (Fig. 1). TM3’seq was developed to fill a current gap in RNA-seq protocols that use late multiplexing (i.e. individual samples are pooled after final PCR amplification); by building up on recent advances in library preparation (lohman 2016, picelli 2014), TM3’seq allows the high-throughput processing of hundreds of individual samples at a fraction of the cost of commercially available kits with equal high-quality performance. With the aim of making TM3’seq truly high-throughput all steps have been streamlined and optimized for 96-well plate processing in liquid-handling platforms without sacrificing library quality.

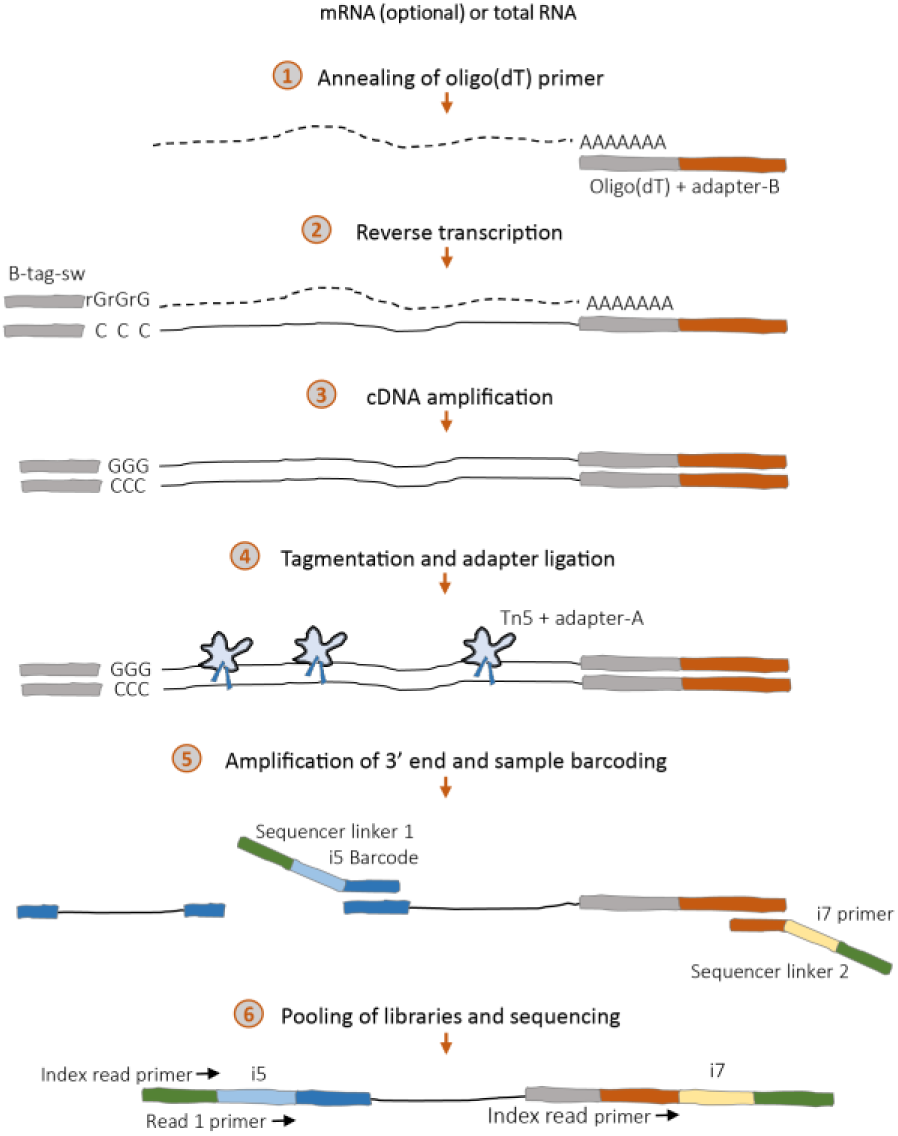

### Performance of TM3’seq in differential expression analysis of human RNA

One of the most common goals of RNA-seq studies is to detect differentially expressed genes between pre-determined groups. Here we have used human blood and adipocyte samples in order to both evaluate the performance of TM3’seq in detecting differentially expressed genes between these two tissues, and to compare such performance to the NEBNext Ultra protocol. A comparison of mapping quality parameters between the methods can be found in Suppl. Fig. 1.

#### Transcriptional profile of blood and adipose tissue

Both methods reliably recover the transcriptional profile of each tissue. In terms of transcript abundance, the correlation between replicates is very high within each method, as wells as between methods (rho > 0.9 for any comparison) (Fig. 2). Regarding the total number of genes detected, there are some tissue-specific differences between methods (Fig 3a). In complex transcriptional profiles like that of adipocytes, both methods recover not only the same number of total genes, but also the same proportion of genes in different abundance groups. However, in transcriptomes dominated by one gene, as is the case of hemoglobin in blood samples, TM3’seq detects slightly fewer genes (∼12%) than NEB. This might be a consequence of the cycles of whole-mRNA amplification performed during cDNA synthesis *in TM3’seq that are not part of the NEBNExt^®^ protocol (See Fig. 1)*.

**Figure 2.**
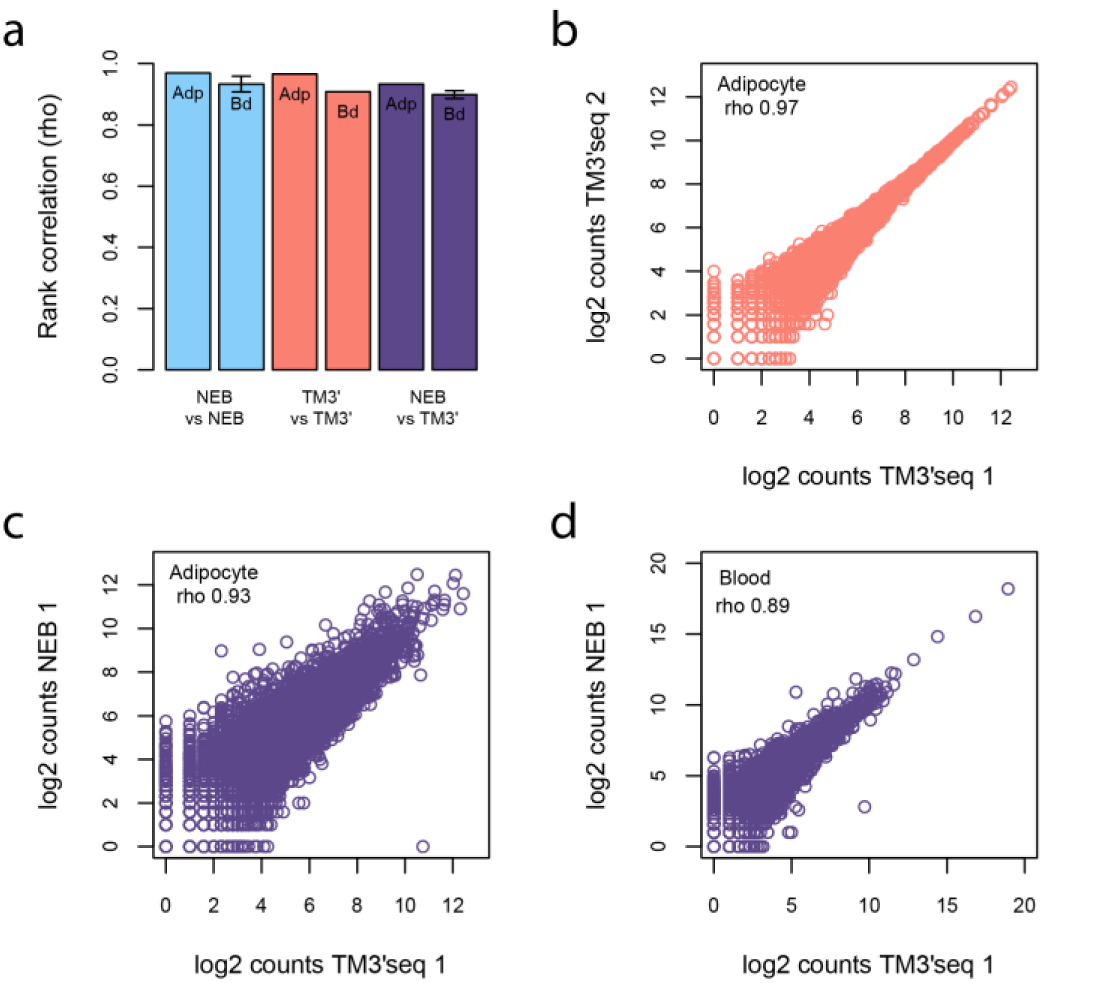
Performance of the TM3’seq method based on the correlation between RNA-seq library replicates. Gene counts per sample were compared between TM3’seq replicates (red), NEB replicates (blue), and between NEB and TM3’seq replicates (purple). Each method has three replicates per tissue (blood – Bd, and adipocytes - Adp). Each replicate corresponds to 1M uniquely mapped RNA-seq reads that were assigned to the set of 20612 protein-coding genes in the GRCh38 assembly of the human genome. TM3’seq libraries were amplified for 12 PCR cycles. Panel (a) shows the average correlation between NEB replicates (n=3), TM3’seq replicates (n=3), and NEB vs TM3’seq samples (n=9). The whiskers indicate two standard deviations from the mean. In most of the groups the standard deviation is too small to be plotted. Panels (b-d) show examples of the correlation between replicates. A comparison between mapping parameters of both methods can be found in Suppl. Fig. 1.

**Figure 3.**
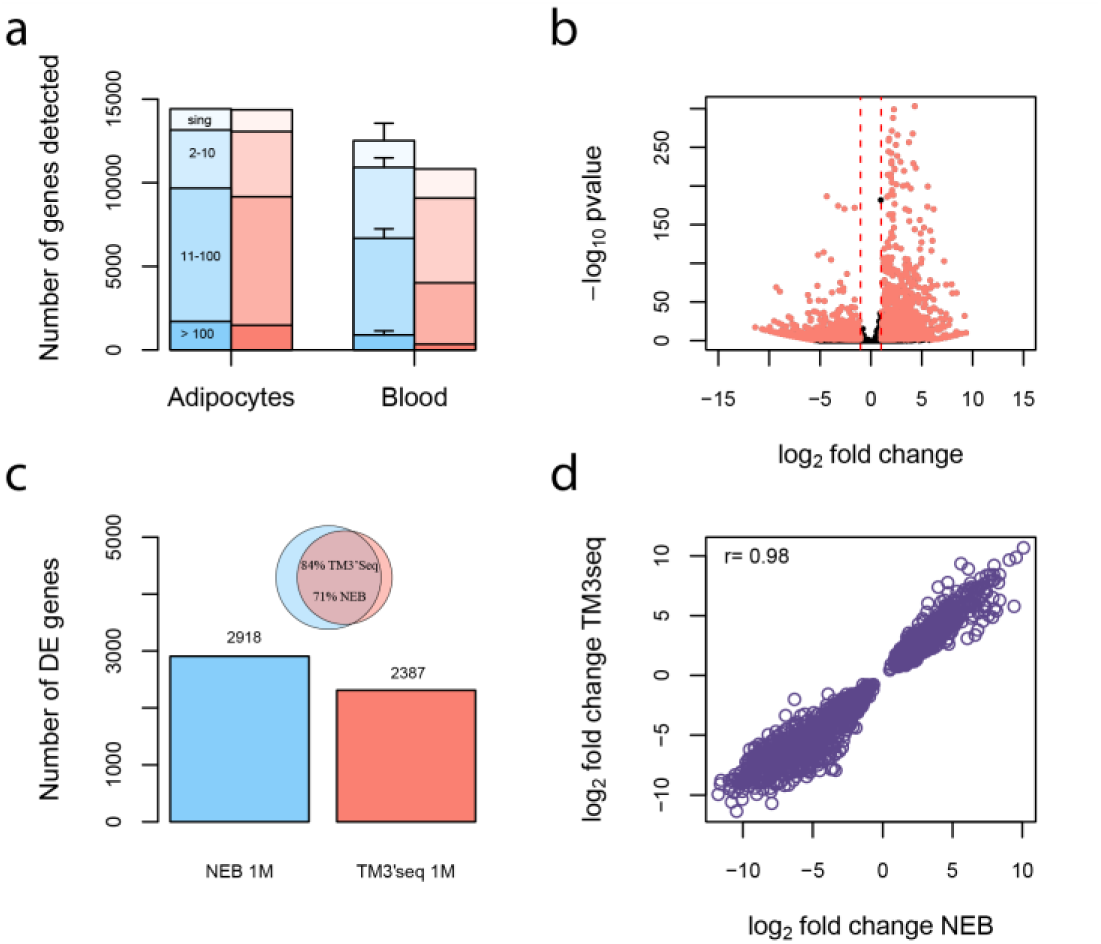
Performance of TM3’seq method based on differential expression analysis between blood and adipocytes. Three replicates per tissue were used. Each replicate corresponds to 1M uniquely mapped RNA-seq reads per sample assigned to the set of 20612 protein-coding genes in the GRCh38 assembly of the human genome. The performance of TM3’seq (red) is compared to NEB@Next Ultra (blue). TM3’seq libraries were amplified for 12 PCR cycles. (a) Number of genes detected with each method. Genes are clustered by abundance: singletons (sing), 2-10 reads, 11-100 reads, and > 100 reads. The average number of genes in three replicates is shown; the whiskers represent two standard deviations; in most samples std is too small to be plotted. (b) Volcano plot showing the significance as a factor of log fold change (lfc) for each gene (dot) in TM3’seq. Significant genes with lfc > 1 are highlighted in red. (c) Differentially expressed genes in each method (Bonferroni p-value <0.05). The inset shows the overlap between differentially expressed genes detected by NEB and TM3’seq at 1M reads. (d) Correlation between the effect sizes (lfc) of the 2082 genes that overlap between TM3’seq and NEB 1M reads.

#### Differential expression analysis

Three replicates per tissue were used to detect differentially expressed genes between blood and adipocytes. With a sequencing depth of 1M reads, TM3’seq is able to detect 2387 differentially expressed genes, while NEB detects 2918 (Fig. 3b). Such difference between methods is small, and natural consequence of the lower number of genes detected in TM3’seq blood samples relative to NEB blood samples (see Fig. 3a). However, and arguably more importantly, the effect size (fold change) of the 2082 overlapping genes is almost identical between methods (r= 0.98, Fig. 3d).

#### Effect of sequencing depth and amplification cycles

In order to explore the effect of sequencing depth in differential expression analyses, we have compared TM3’seq samples with 1M, 2M, and 3M uniquely mapped reads (Suppl. Fig. 2). The number of detected genes, and the number of differentially expressed genes increases with depth of sequencing, as expected. In adipocytes, 2M- and 3M-reads samples detect 6% and 8% more genes than 1M sample, respectively (10% and 16% in blood). However, it should be noted that although this gain in information seems modest regarding the total number of transcripts, it becomes more important in high abundance transcripts (>100 reads) (Suppl. Fig. 2a). The number of transcripts with 100 or more reads is 133% and 242% higher in 2M- and 3M-reads samples than in 1M-reads samples, respectively (125% and 265% in blood). The increase in the number of transcripts with moderate to high gene counts results in higher power to detect differentially expressed genes; samples with 2M and 3M reads detect 52% and 84% more differentially expressed genes than samples with 1M reads (Suppl. Fig. 2b).

Contrary to sequencing depth, the number of library amplification cycles does not substantially affect the number of genes detected, nor the ability to call differentially expressed genes (Suppl. Fig. 2c-d). Consequently, if desired or needed, the library concentration can be increased by increasing the final number of PCR cycles without affecting the biological information contained in the sample.

### Performance of TM3’seq based on ERCC probes

ERCC probes are poly-A RNA probes that span a known range of concentrations and lengths. Given such known concentrations, ERCC probes were used to compare the technical performance of NEBNext^®^ and TM3’seq methods. Both methods show high correlation between replicates (rank correlation rho > 0.95) (Suppl. Fig. 3a), and both have limitations in detecting lowly expressed probes (Suppl. Fig 3b). Overall, technical performance is very similar between methods, and it is robust even when sequencing depth is as low as 100k reads per sample.

### Analysis of head and body transcriptomes of Drosophila melanogaster

As a proof of principle, we have used our newly developed TM3’seq protocol to analyze the transcriptomes of 96 outbred *Drosophila melanogaster* flies. We have processed heads and bodies separately for each individual fly. Our results show that TM3’seq works reliably for high-throughput processing of low-input samples (single fly head) and moderate-input samples (single fly body), allowing mainstream comparison of transcriptomes (Fig. 4).

**Figure 4.**
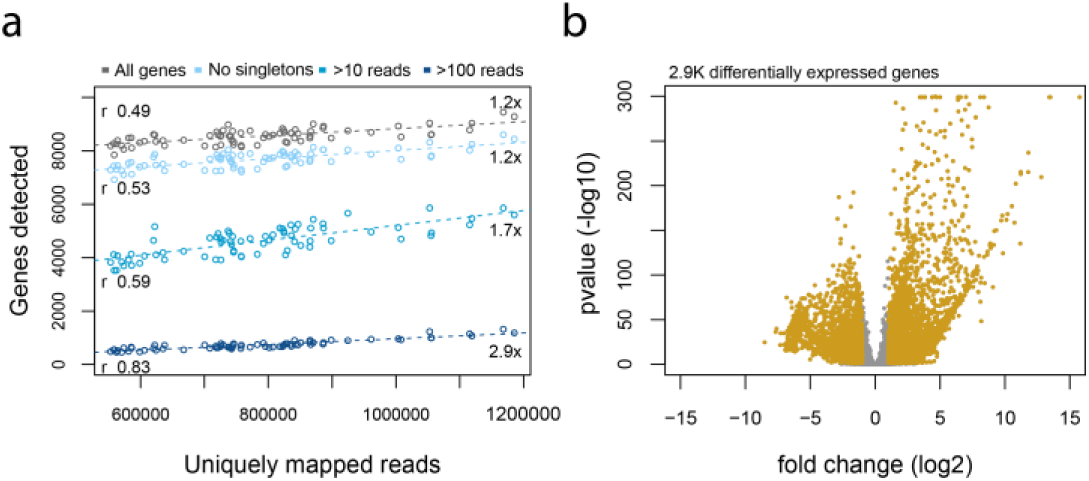
Transcriptome analysis of *Drosophila melanogaster* using TM3’seq. 48 flies were profiled for head and body transcriptomes. Panel (a) shows the relationship between the number of genes detected given depth of sequencing for each individual library. The strength of the correlation is shown in the left side of the panel, and the increase in number of genes between samples with the highest (1.2M) and lowest (500K) sequencing depth is shown in the left. Each dot represents an individual fly transcriptome. The different lines represent a different category of genes determined by their abundance. (b) Volcano plot showing the results of differential gene expression analysis between body and head. Each dot represents a gene. Yellow dots are significant genes (Bonferroni p-value <0.05) with a log fold change > 1. Positive fold change indicates genes overexpressed in heads relative to bodies.

## Discussion

The cost and throughput of library preparation are current limiting factors for the scalability of RNA-seq experiments. Consequently, many attempts are being made to reduce reagent costs in a way that keeps up with the reduction in sequencing costs. The most successful approach, early multiplexing of individual samples, came from the field of single-cell transcriptomics (Macosko et al. 2015, Hashimshony et al. 2016). However, for some experimental designs, early multiplexing is not feasible, and current protocols using late multiplexing have not been able to match the throughput and low costs of early multiplexing approaches. The method presented here, TM3’seq, aims to fill such gap by offering a high-quality, high-throughput, and low-cost RNA-seq library preparation method that uses late multiplexing.

### Cost and High-throughput

The current reagent cost of RNA-seq library preparation using commercial kits is very high: $35 - $44 per sample (NEBNext^®^Ultra kit). Our protocol offers a much-needed reduction not only in cost per sample, but also in hands-on time. The reagent cost per sample using TM3’seq is $1.5, cheaper than any of the early multiplexing approaches (e.g. SCRB-seq ∼$2.2, BRB-seq ∼$2.2), and one person can process 96 samples in 5-6 hours, with only 2-3 hours of hands-on time.

To further reduce sequencing costs, TM3’seq implements the dual-indexing approach from Illumina^®^ in a way that both indexes can be read using single-end sequencing. This allows high multiplexing of samples while eliminating the requirement for pair-end sequencing of early-multiplexing approaches.

Beyond reagent and sequencing costs, another limiting step in terms of cost and scalability of RNA-seq studies is RNA extraction. This is traditionally thought of as independent from library preparation optimizations, however in order to make TM3’seq feasible for hundreds or thousands of samples, we have also optimized the mRNA extraction step to work with low-input samples (e.g. single *Drosophila melanogaster* heads) and in a cost-efficient manner following Kumar et al. (2012). The cost per sample is ∼$1.7, and samples can be processed in 96-well plates; this is ∼5 times cheaper than the column-based PicoPure™ RNA Isolation kit (ThermoFisher) that only allows parallel procession of a few samples. It should be noted that currently our protocol is not optimized to work with ultra-low input (e.g. single cell RNA amount) but could be adapted to such conditions by optimizing the number of cDNA amplification cycles. To further make the process as high-throughput as possible, all our protocols were optimized for 96-well plates from the first step (mRNA extraction) to the last (PCR amplification of libraries) (Suppl. File 1, Suppl. File 2, and webpagelink).

### Preservation of sample individuality

Other approaches to reduce the cost and increase the throughput of RNA-seq library preparation come from the single-cell sequencing field. Most of them rely on the barcoding and pooling of samples before second strand cDNA synthesis; the resulting pool is then treated as a single sample for the remaining steps of the protocol (Macosko et al. 2015, Hashimshony et al. 2016, Alpern et al. 2019). This early multiplexing approach means that the identity of each sample is only recovered after the sequencing results are analyzed. This feature is not ideal for most studies that focus on samples other than single cells (e.g. tissues, organism). For such experiments, it is indeed desirable or necessary to preserve the identity of each individual library for future follow up studies.

Our protocol does not pool the samples before reverse transcription, but treats them individually along each step of the protocol, resulting in individual libraries for each sample. The barcodes are added during the last PCR amplification step, and the multiplexing is done right before sequencing. This feature allows the re-sequencing of specific samples for either, deeper exploration of interesting samples, or to increase read depth for shallowly covered samples. Importantly, the TM3’seq cost per sample is lower than any of the early multiplexing protocols (e.g. SCRB-seq ∼$2.2, BRB-seq ∼$2.2), and the processing time is comparable.

Another caveat of early multiplexing approaches is that any between-sample variation in RNA input amount will result in high variation in sequencing coverage per sample; therefore, the precise quantification of RNA input previous to sample pooling and reverse transcription is mandatory (Alpern et al. 2019). By using late multiplexing, the TM3’seq protocol allows the pooling of individual libraries in the desired proportion before sequencing, assuring that reads will be distributed as desired by the researcher.

### TM3’seq performance

We have simplified the TM3’seq protocol as much as possible to make it truly high-throughput and inexpensive without sacrificing the quality of the resulting RNA-seq libraries. This is reflected in the high technical and biological performance of TM3’seq, comparable to the high quality of the commercial gold standard in the field, NEBNext^®^ Ultra. Both methods show equivalent performance, not only in the number of genes detected, but also in the limits of RNA molecule detection set by the length and concentration of known mRNA probes (ERCC probes).

The validation of TM3’seq using diverse human tissues, as well as *Drosophila* body parts, and artificial RNA molecules (ERCC probes) shows the reliability of TM3’seq in a wide spectrum of transcriptome profiles. The optimization of mRNA extraction for low input tissue as well as the high-throughput of our method was evaluated with *Drosophila melanogaster* single heads and bodies. High quality libraries were obtained from as little as 10ng of mRNA (one *Drosophila* head).

We did not include unique molecular identifiers –UMI-in this protocol given that accounting for PCR duplicates in RNA-seq experiments does little to the power to detect differentially expressed genes (Alpern et al. 2019). Furthermore, UMIs are rarely used in transcriptomic studies beyond single-cell sequencing. However, if the experimental design requires unique molecular identifiers, they can be custom-added to the standard Illumina i5 (see Suppl. File. 3) or i7 oligos. Adding UMI to the i5 oligo still allows for single-end sequencing, therefore preserving the TM3’seq reduction in sequencing costs.

## Conclusion

Our results indicate that TM3’seq is a reliable, inexpensive, and high-throughput method to prepare RNA-seq libraries. The protocol is straightforward and has been optimized to work in a 96-well plate set up; it therefore can be easily implemented in any laboratory for the high-throughput and cost-effective processing of large scale RNA-seq projects. To further facilitate the analysis of the data, we also make available a pipeline that allows the researcher to go from raw FASTQ files to gene counts files ready to be analyzed in standard differential expression programs like DEseq2 (Love et al. 2014), or used as input for eQTL mapping in MatrixEQTL (Shabalin 2012). With TM3’seq RNA-seq, experiments can be significantly scaled up, enabling large sample sizes necessary to address questions in the fields of quantitative genetics, phenotypic prediction, and systems genetics.

## Supporting information

Supplemental File 1 - Library preparation protocol

Supplemental File 2 - mRNA extraction protocol

## Acknowledgments

We thank Jen Grenier, Jethary Rader, and the Ayroles lab for comments on early drafts of this manuscript. We also thank Lance Parsons for help setting up the data analysis pipeline and the website. We especially thank Anett Schmittfull for her help and input while optimizing this protocol. This study was funded by NIH R35 GM124881 and R01ES029929 to JFA. LFP is funded by a Long-Term Postdoctoral Fellowship from the Human Frontiers Science Program.

