## Supplemental File 3 for "TM3’seq: a tagmentation-mediated 3’ sequencing approach for improving scalability of RNA-seq experiments"

**Supplementary File 3.** Incorporation of UMIs into the TM3’seq protocol

Unique molecular identifiers have been extensively used in single-cell transcriptomics to account for the amplification bias resulting from the high number of whole-transcriptome amplification cycles. Given that other kinds of RNAseq experiments start with larger amounts of RNA input, UMIs are not widely used outside of single-cell transcriptomics field. However, since it might be desirable to include UMI in certain types of experiments, here we show how to implement UMIs into the TM3’seq protocol.

We suggest that UMI barcodes be added to the i5 oligo given that his allows to preserve the TM3’seq single-end sequencing property. The UMIs can be also added to i7 oligo, but this will require pair-end sequencing.

To incorporate UMIs in the i5 oligo, a sequence of 8 random nucleotides is added to the standard i5idx Illumina barcode, where **N** indicates the i5 barcode, and **n** the UMI:

AATGATACGGCGACCACCGAGATCTACAC**NNNNNNNNnnnnnnnn**TCGTCGGCAGCGTC

The TM3’seq protocol can be followed in the same way as described in Suppl. File 1, with an additional step cleaning step at the final library amplification step:

- **Final library amplification**:

In this final step, 10ul of OneTaq HS Quick-Load 2x (NEB, #M0486L), 1ul i5 primer (i5 barcode + UMI) 1uM, 1µl i7 primer 1µM, and 7µl of water are used to amplify 1µl of the tagmentation reaction following the program: 68°C 3min, 95°C 30sec, [95°C 10sec, 55°C 30sec, 68°C 30sec] ***3 cycles**, 68°C 5min. These three cycles allow the incorporation of unique i5 primers (and therefore unique UMI) to each RNA molecule. To avoid the unspecific amplification of RNA molecules by free i5 primers, the samples are pooled and cleaned using a ratio of 1x Agencourt AMPure XP beads (Beckman Coulter, #A63881). The pool of samples is then amplified for extra 12cycles (or how many are desired) using i5 primes that do not contain UMI barcodes following the program: 68°C 3min, 95°C 30sec, [95°C 10sec, 55°C 30sec, 68°C 30sec] *15 **cycles**, 68°C 5min.
