## Supplemental Table 1 for "TM3’seq: a tagmentation-mediated 3’ sequencing approach for improving scalability of RNA-seq experiments"

**Supplementary Table 1.** Oligonucleotides used in TM3’seq protocol. All oligos were synthesized using standard desalting by Integrated DNA Technologies – IDT.

| **Oligo name** | **Oligo sequence** |
| --- | --- |
| Tn5ME-B-30T | 5’-GTCTCGTGGGCTCGGAGATGTGTATAAGAGACAGTTTTTTTTTTTTTTTTTTTTTTTTTTTTTTV-3' |
| B-tag-sw | /5Biosg/ACCCCATGGGGCTACACGACGCTCTTCCGATCTrGrGrG |
| Adapter-A (Illumina) | 5’- TCGTCGGCAGCGTCAGATGTGTATAAGAGACAG-3’ |
| Tn5MErev (Picelli 2014) | 5’-[phos]CTGTCTCTTATACACATCT-3’ |
| i5 (Illumina Idx5) | AATGATACGGCGACCACCGAGATCTACACNNNNNNNN*TCGTCGGCAGCGTC |
| i7 (Illumina Idx7) | CAAGCAGAAGACGGCATACGAGATNNNNNNNN*GTCTCGTGGGCTCGG |

*N, indicates i7 or i5 barcodes. Standard Illumina barcodes were used.
