## Supplemental Figures for "TM3’seq: a tagmentation-mediated 3’ sequencing approach for improving scalability of RNA-seq experiments"

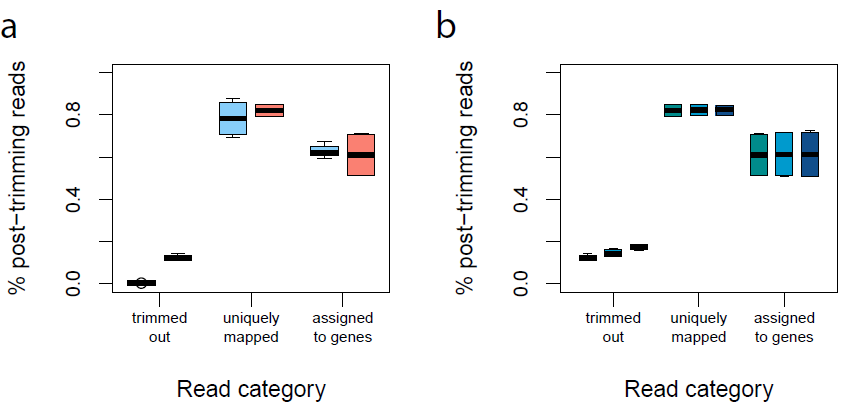

**Supplementary Figure 1. Mapping parameters.** (a) NEB (blue) and TM3’seq (red) libraries (six samples per method), and (b) TM3’seq libraries amplified for 12, 15, and 18 cycles were compared in terms of the reads removed due to sequencing adapters, the percentage of uniquely mapped reads after trimming, and the percentage of such reads that were assigned to protein coding genes.

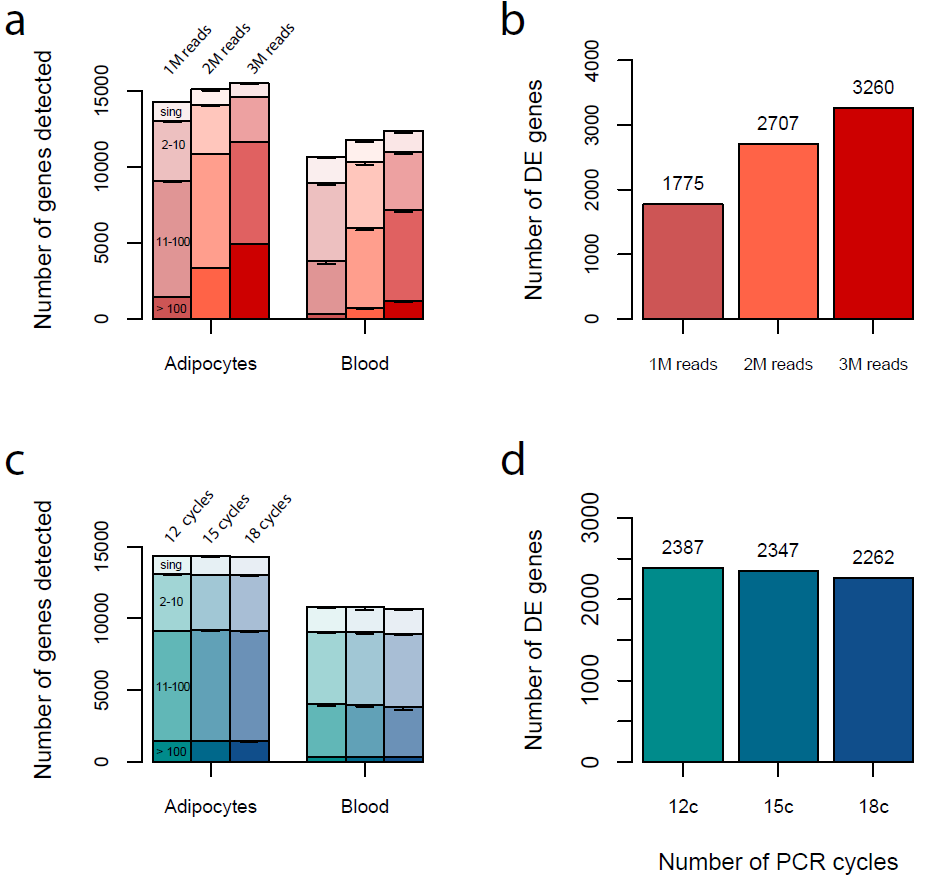

**Supplementary Figure 2. Effect of sequencing depth and amplification cycles on the number of genes detected and the number of genes differentially expressed between blood and adipocytes using TM3’seq.** (a,b) Libraries with 1M, 2M, and 3M uniquely mapped RNA-seq reads were amplified for 18 cycles. Two adipocyte and three blood replicates were used. (c,d) Libraries were amplified for 12, 14, and 18 cycles, each sample was down sampled to one million uniquely mapped reads. Each tissue has three technical replicates. The average number of genes detected is shown in (a) and (c) Genes are clustered by abundance: singletons (sing), 2-10 reads, 11-100 reads, and more than 100 reads. Whiskers represent two standard deviations. For some bins, std is too small to be plotted. The number of differentially expressed genes (Bonferroni p-value <0.05) is shown in (b) and (d). The differentially expressed genes identified by each number of cycles are very similar, with the overlap between cycles ranging from 88% to 98%.

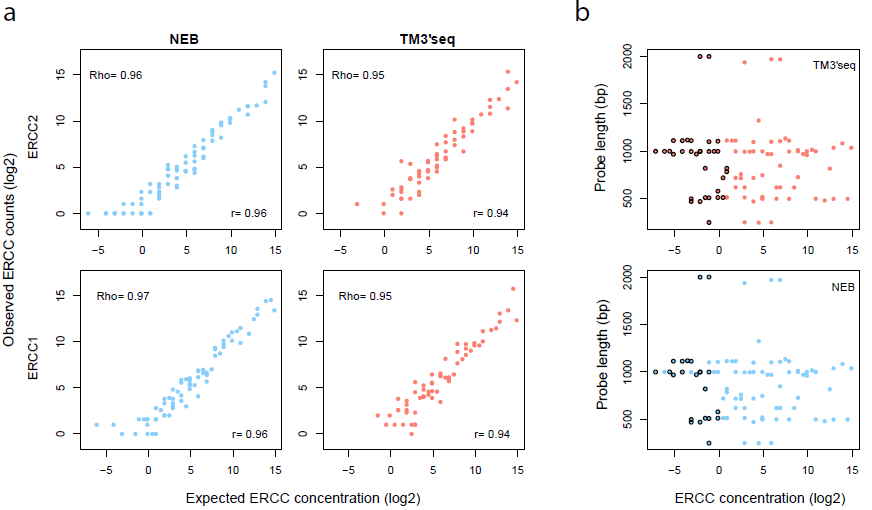

**Supplementary Figure 3. Technical performance of TM3’seq and NEB methods based on detection of ERCC probes.** (a) Expected vs observed ERCC amounts. Pearson (r2) and Spearman rank (rho) correlations are shown; each value corresponds to the average of two replicates. (b) Detection performance of ERCC probes given the length and the concentration of the probe. Each dot represents an ERCC probe, dots with black border represent the probes that were not detected in any of the four samples per method, or that have a mean expression <1 based on the output of ERCC-Dashboard (Munro, Lund et al. 2014).
