## Supplemental File 1 - Library preparation protocol for "TM3’seq: a tagmentation-mediated 3’ sequencing approach for improving scalability of RNA-seq experiments"

**Supplemental File 1.** TM3’seq protocol

**Reagents and materials:**

**(a)**	SMARTScribe ™ RT (Takara, #639538)

**(b)**	dNTPs (NEB, #N0447S)

**(c)**	DTT (Takara, #639538)

**(d)**	5x First-Strand buffer (Takara, #639538)

**(e)**	OneTaq HS Quick-load 2x (NEB, #M0486L

**(f)**	Re-association buffer

**(g)**	Re-association buffer:glycerol

**(h)**	TAPS buffer 5x

**(i)**	SDS (Promega, #V6551)

**(j)**	Agencourt AMPure XP beads (Beckman Coulter, #A63881)

**Buffers:**

**(a)**	Re-association buffer [store at room T] 10mM Tris pH 8.0, 50mM NaCl, 1mM EDTA.

**(b)**	Re-association buffer:glycerol 1:1 [store at room T], mix one volume of re-association buffer and one volume of glycerol.

**(c)**	TAPS buffer 5x pH 8.5 from Picelli, Faridani et al. (2014) [store at 4°C], 50mM TAPS, 25mM MgCl_2_, 50% v/v DMF.

**Oligos:** See supplementary table 1

**Equipment: T**hermal cycler

- **First strand cDNA synthesis**

1.	Pre-heat a thermal cycler to 65°C.

2.	Start with a minimum of 200ng total RNA or 10-20ng of mRNA in 10µl.

[*These are the input amounts for which TM3’seq has been optimized, however, we have tested this protocol with input amount ranging from 100ng to 1000ng of total RNA without any major differences in performance*]

3.	Add 1µl of 0.83µM Tn5ME-B-30T oligo to each RNA sample and incubate at 65°C for 3min. Then, place immediately on ice to prevent the dissociation of oligo-dT oligos from the polyadenylated tail of RNA molecules.

[*This is the step when the Tn5ME-B-30T oligo binds to the polyA tail of mRNA molecules. Besides a polyT tail, this oligo contains the complementary sequence of the i7 primer that will be used to both, add the sequencing adapter and i7 index to each sample in the final library amplification and barcoding step*]

4.	Prepare reverse transcription (RT) master mix for each sample. Include reagents for one or two extra samples -depending on how many samples are being processed- to account for pipetting errors.

| **Reagent** | **Volume (µl)** |
| --- | --- |
| dNTP (10mM) | 1 |
| DTT (0.1M) | 2 |
| 5x 1st-strand buffer | 4 |
| B-tag-sw (0.83µM) | 1 |
| SMARTScribe RT | 1 |

5.	Add 9µl of RT master mix to each RNA sample for a total volume of 20µl and mix thoroughly by pipetting up and down. Incubate in a thermocycler for 1h at 42°C for the reverse transcription to occur, and 15min at 70°C to inactivate the reverse transcriptase.

- **cDNA synthesis** **and amplification**

6.	Prepare cDNA amplification master mix for each sample. Include reagents for one or two extra samples -depending on how many samples are being processed - to account for pipetting errors.

| **Reagent** | **Volume (µl)** |
| --- | --- |
| OneTaq HS Quick Load | 7.5 |
| Nuclease-free water | 2.5 |

7.	Mix 10µl of cDNA master mix with 5µl of RT reaction, mix thoroughly by pipetting up and down until solution looks homogeneous. Or, vortex the tubes and spin them down to collect the all the liquid.

8.	Perform three rounds of whole cDNA amplification using the following program in a thermal cycler:

| **Step** | **Temperature** | **Time** |
| --- | --- | --- |
| 1 | 68°C | 3min |
| 2 | 95°C | 30sec |
| 3  (3 cycles) | 95°C | 10sec |
|  | 55°C | 30sec |
|  | 68°C | 3min |
| 4 | 68°C | 5min |
| 5 | 4°C | Hold |

- **Anneal the adapters that will be loaded into the Tn5 transposase**

***Note:*** *This step can be done once, and the annealed adapters can be placed at 4°C for long-term storage. The annealed adapters stored at 4°C can be directly used in step 10 of this protocol.*

9.	Anneal the adapter-A forward and reverse oligos by mixing 10µl adapter-A oligo (100µM), 10µl Tn5MErev oligo (100µM), and 80µl of re-association buffer. To anneal the oligos run the following program in the thermal cycler.

| **Step** | **Temperature** | **Time** |
| --- | --- | --- |
| 1 | 95°C | 10min |
| 2 | 90°C | 1min |
| 3  (60 cycles) | Reduce the temperature by 1°C every cycle | 1min per cycle |
| 4 | 4°C | Hold |

[*The adapter-A oligo is the free-end adapter that will be ligated to the cDNA after the Tn5 transposase fragments the cDNA.* *This oligo contains the complementary sequence of the i5 primer that will be used to both, add the sequencing adapter and i5 index to each sample in the final library amplification and barcoding step*]

- **Load the Tn5 with adapter-A**

10.	Load the annealed adapter-A into the Tn5 transposase by mixing one volume of homebrew Tn5 and one volume of 1µM annealed adapter-A oligo. Incubate the mix in a thermal cycler with pre-heated lid for 30min at 37°C.

11.	Make a 1:7 dilution of the loaded Tn5 using a 1:1 mix of re-association buffer and glycerol.

[*Steps 10 and 11 are highly variable depending on the specific Tn5 being used and the desired library size, where small average size libraries can be obtained by increasing Tn5 concentration. Given that more and more laboratories are purifying their own Tn5, each one will have to adjust the concentration of Tn5 based on their needs and the specific properties of their Tn5. However, we have found that the yield of the final library increases when Tn5 is loaded with adapter-A in the absence of any buffer, and therefore we suggest that the dilution of Tn5 be done after the adapter-A has been loaded*]

- **cDNA tagmentation**

12.	Prepare the tagmentation master mix for each sample. Include reagents for one or two extra samples -depending on how many samples are being processed - to account for pipetting errors.

| **Reagent** | **Volume (µl)** |
| --- | --- |
| Loaded Tn5 | 1 |
| TAPS buffer 5x | 4 |
| Nuclease-free water | 5 |

13.	Mix 10µl of tagmentation master mix with 5µl of cDNA (from step 8). Mix thoroughly by pipetting up and down until the solution looks homogeneous, and incubate the mix in thermal cycler for 7min at 55°C.

14.	Dissociate the Tn5 that remains bound to cDNA molecules by adding 3.5µl of 0.2% SDS to each sample. Mix very well by pipetting up and down. Incubate the mix in a thermal cycler for 7min at 55°C

[*It is critical that the 0.2% SDS dilution is prepared fresh from the stock every time a tagmentation reaction is performed. Old, diluted SDS does not efficiently remove the Tn5 from the cDNA and this will result in lack of library amplification. It is also important not to increase the volume of SDS added to the reaction because this will inhibit the following PCR amplification step*]

- **Library amplification and sample barcoding**

15.	Prepare the PCR master mix for each sample. Include reagents for one or two extra samples -depending on how many samples are being processed- to account for pipetting errors.

| **Reagent** | **Volume (µl)** |
| --- | --- |
| i5 primer (1µM) | 1 |
| OneTaq HS Quick Load | 10 |
| Nuclease-free water | 7 |

[*Here, we use the i5 primer as the “batch” or “plate” barcode, so the same i5 is added to each sample. It can be used to label each 96-well plate being processed. In the next step, i7 primers are used as “sample” barcodes, and therefore a unique i7 per sample is used. i7 will therefore barcode each well, and i5 will barcode each plate/batch. This allows high multiplexing of samples that will be sequenced together. The way i5 and i7 primers are used to barcode samples can be adjusted depending on the experimental set up*]

16.	Mix 1µl of each tagmented cDNA sample (from step 14) with 18µl of PCR master mix and 1µl of i7 primer (1µM). Mix thoroughly by pipetting up and down.

[*The i7 primer should be unique for each sample, see comment on step 15. We do not recommend changing the volumes of this particular reaction because higher amounts of SDS easily inhibit the PCR reaction*]

17.	Amplify the samples using the following program in a thermal cycler:

| **Step** | **Temperature** | **Time** |
| --- | --- | --- |
| 1 | 68°C | 3min |
| 2 | 95°C | 30sec |
| 3  (18 cycles) | 95°C | 10sec |
|  | 55°C | 30sec |
|  | 68°C | 30sec |
| 4 | 68°C | 5min |
| 5 | 4°C | Hold |

[*The number of PCR cycles can be adjusted depending on the amount of RNA input and desired yield. We use 18 cycles when working with single Drosophila head, but we have obtained enough yield starting from 12 cycles. The number of cycles at this step does not affect the quality of the final library (see Fig3b-d)*]

- **Library cleaning and pooling**

18.	At this point, each individual library has a unique barcode, and therefore they can be pooled and cleaned, or they can be cleaned individually. Perform a double-sided clean up (0.6x-1x ratio) using Agencourt AMPure XP beads to reduce the amount of primer dimmers that will be carried over for sequencing.

19.	Quantify the concentration of the libraries by Qubit and check the quality using Bioanalyzer or TapeStation. A normally distributed library with a peak ranging from 300-500 is ideal (Fig. 1a). If a primer dimer peak is still visible at ~150bp we recommend cleaning the library again before sequencing using a bead ratio of 1x.

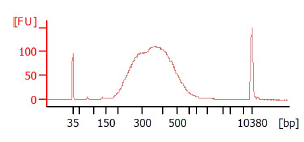

**Figure 1**. Bioanalyzer trace of a pool of 96 single head *Drosophila melanogaster* libraries processed in a 96-well plate following the TM3’seq protocol. No primer-dimer peak visible at 150bp.

**--**
