## Supplemental File 2 - mRNA extraction protocol for "TM3’seq: a tagmentation-mediated 3’ sequencing approach for improving scalability of RNA-seq experiments"

**Supplemental File 2.** mRNA extraction procedure

- **mRNA extraction** [*This is an* o*ptional step given that total RNA can be used as input material for TM3’seq. RNA extraction was chosen instead of total RNA isolation because it can be implemented in 96-well plates and liquid-handling platforms to increase throughput. The mRNA isolation protocol is based on* *Kumar, Ichihashi et al. (2012)*]

**Reagents and materials:**

**(a)**	Dynabeads™ mRNA DIRECT™ Purification Kit (ThermoFisher, #61012). This kit includes oligo(dT) beads, Lysis/Binding buffer –LBB, washing buffer A and B, and Tris-HCl.

**(b)**	Homebrew LBB buffer. In order to reduce costs when processing many samples, we suggest using homebrew LBB buffer for the washing steps and using the kit LBB for tissue homogenization.

**(c)**	Homebrew low-salt buffer – LSB.

**(d)**	2.8mm Stainless steel grinding beads (OPS diagnostics, #089-5000-11).

**(e)**	96-well PCR plates (or PCR strips if processing less samples) with PCR strip lids

[*Important: make sure to use lids that prevent leaks during sample homogenization*].

**Buffers:**

**(a)**	Homebrew LBB [store at 4°C], 100mM Tris-HCl pH 7.5, 500mM LiCl, 10mM EDTA pH 8.0, 1% SDS, 5mM DTT.

**(b)**	Homebrew LSB [store at 4°C], 20mM Tris-HCl pH 7.5, 150mM NaCl, 1mM EDTA.

**Equipment:**

**(a)**	Any homogenizer that can handle 96-well plates, we used Talboys High Throughput Homogenizer (#930145).

**(b)**	96-well plate magnet (Invitrogen™, Magnetic Stand-96, AM10027).

- **Preparing the buffers**

1.	Take all buffers and Dynabeads out of their 4°C storage for at least 30min before starting the protocol. Mix thoroughly to make sure any precipitate that might have formed is dissolved.

2	Add β-mercaptoethanol and Antifoam A to the LBB buffer before homogenization of the tissue. LBB+ = 10ml LBB, 150µl Antifoam A, 50µl β-mercaptoethanol.

[*If working with LBB from the Dynabeads kit and homebrew LBB, add β-mercaptoethanol and Antifoam A to both. LBB makes a lot of foam during homogenization. When working with low tissue input like Drosophila heads, any amount of RNA lost to the foam will determine whether the library preparation will be successful or not. In our experience, the modifications made by Kumar, Ichihashi et al. (2012) to LBB buffer to reduce foam were determinant for the successful library preparation from a single fly head.*]

- **Homogenizing the tissue**

3.	Add one grinding bead to each well of the 96-well plate containing the samples. Keep the plate on a pre-chilled rack.

4.	Add 100µl of LBB+ buffer to each well. Put the PCR strip lids on and place in a homogenizer for 5-10min, max speed.

[*This volume was used for single Drosophila head and single body, but it can be increased if working with larger input samples, however, the PCR tubes will need to be replaced by 1.5ml tubes. If this volume is modified, we recommend modifying the volume used in the other washing steps*]

6.	Keep at room temperature for at least 5min, this will give time for lysis to be completed while the foam goes down.

7.	Centrifuge for 10 minutes to get rid of the rest of the foam. While performing this step, prepare the Dynabeads (step 8). At this point, the lysate can be transferred to a new PCR plate and stored at -80°C if necessary.

- **Preparing the oligo-dT beads**

8.	While samples are being centrifuged (step 7), re-suspend Dynabeads thoroughly before use by turning the bottle upside down repeatedly.

9.	Transfer 20µl beads for 100µl lysate into each well of a new 96-well plate and place it on the magnet. Wait 2-5mins or until the suspension is clear, remove the supernatant without disturbing the beads.

[*This volume of beads was optimized to get as much mRNA from a single Drosophila head at the lowest possible cost; it was also used to process single fly bodies. However, if working with larger samples, this volume can be increased to recover more mRNA*]

11.	Remove the plate from the magnet. Add 100µl of LBB+ buffer to each well, and wash the beads by pipetting up and down until the solution looks homogeneous.

12.	Place the plate on the magnet, wait until suspension is clear, and remove the LBB+ buffer without disturbing the beads. Remove the plate from the magnet.

- **mRNA extraction**

13.	Add the sample lysate (from step 7) to the plate with beads. Transfer as much lysate as possible without taking the debris pellet.

14.	Re-suspend the beads by pipetting up and down several times until the solution looks homogeneous. Incubate at room temperature for 10min using a plate mixer if available. During this step, the polyA tail of RNA molecules will bind to the oligo-dT beads.

[*If final mRNA yield is lower than expected, the incubation time can be increased; we didn’t see any significant improvement when increasing incubation time, but it might become relevant when using higher RNA input*].

15.	Place the plate on the magnet, and wait 2-5min, or until the solution is clear. Remove the supernatant without disturbing the beads. At this step, the mRNA is already bound to the beads. Remove plate from the magnet.

16.	Add 100µl of Washing Buffer A to each well and pipet up and down several times until the beads are totally re-suspended. Place the plate on the magnet, wait until solution is clear and remove the buffer. Be careful not to disturb the beads. Remove plate from magnet.

17.	Add 100µl of Washing Buffer B to each well, and perform the washing step as described in 16.

18.	Add 100µl of LSB buffer to each well and proceed as in 16.

19.	Add 11µl of 10mM Tris-HCl to each well and mix thoroughly to re-suspend the beads. Incubate the plate at 80°C for 2mins to elute the mRNA from the beads. Place the plate immediately on the magnet and wait until the solution is clear. Remove 10µl of supernatant that now contains the mRNA and transfer to a new plate. This mRNA is ready to be used as input for library preparation.

[*To prevent mRNA from binding back to the oligo-dT beads, the plate can be put on ice immediately after the 80°C incubation. This is a critical step if for any reason the plate cannot be placed on the magnet immediately after incubation*]
